# A data-driven approach for enhancing forest productivity by accounting for indirect genetic effects

**DOI:** 10.1101/2023.04.14.536978

**Authors:** Filipe M. Ferreira, Saulo F. S. Chaves, Leonardo L. Bhering, Rodrigo S. Alves, Elizabete K. Takahashi, Marcos D. V. Resende, João E. Souza, Salvador A. Gezan, José M. S. Viana, Samuel B. Fernandes, Kaio O. G. Dias

## Abstract

Maintaining the past decades current genetic gains for tree species is a challenging task for foresters and tree breeders due to biotic and abiotic factors. Planting a mixture of genotypes or clonal composites can be an alternative to increase the phytosanitary security and yield of forest plantations. These clonal composites are more complex than monocultures due to inter-genotypic competition and indirect genetic effects that can affect the total heritable variation. This study aims to understand how indirect genetic effects can impact the response to selection and how the stand composition can be used to explore these effects and enhance forest yield. We used a clonally trial of *Eucalyptus urophylla* × *Eucalyptus grandis* hybrids in a randomized complete block design with 24 replications, containing a single tree per plot evaluated for mean annual increment at 3 and 6 years. We focus on partitioning the genetic variation of trees into direct and indirect genetic effects based on competition intensity factors. We identified clones as aggressive, homeostatic, and sensitive based on the magnitude of indirect genetic effects. By accounting for indirect genetic effects, for mean annual increment, the total heritability decreased 39 and 44% for 3 and 6 years, respectively. We proposed a workflow that uses the direct and indirect genetic effect to predict the mean value of clonal composite combinations and to select the one with highest yield. Our methodology accounted for spatial variability and interplot competition that can contribute to the total heritable variance and response to selection in forest trials. Based on the models evaluated, the clones are easily classified according to their deviation from the indirect genetic effects mean. Also, we extract useful information to predict different clonal composites compositions, their expected average performance, and define the best recommended combination to be planted in large scale.

## 1. Introduction

In the past decades, there was great increase in planted forest productivity due to high genetic gains. Nowadays, the demands are regard to increase genetic gains, which is still low compared to the observed potential. Future challenges can be related to maintain current growth rates. Projections for the coming years suggest that changes in temperature and precipitation patterns will increase the vulnerability of forests to drought (Choat et al., 2012), pests and diseases (Trumbore et al., 2015), and forest fires (Canadell et al., 2021). These aggravating productivity reducer factors are forcing changes into breeding strategies. Forest tree breeders are concerned with keeping populations with sufficient genetic diversity, capable of sustaining the harmful influence of biotic and abiotic stresses (Rezende et al., 2019). Forests with greater diversity are expected to be more resilient to climate changes consequences (Morin et al., 2018). The maintenance of wood productivity depends on forest composition (Coomes et al., 2014), stand structural homogeneity (Soares et al., 2016) and the choice of individuals with stable phenotypic response (Nicotra et al., 2010).

Recent evidence has shown the potential of cultivating a mixtures of improved clones, also called clonal composites (CC), to increase productivity (Rezende et al., 2019) and to mitigate the effects of genotypes by environments interaction (Oliveira et al., 2023). Comparing CC with monoclonal planting, Rezende et al. (2019) found a 9.8 and 6.3 % increase in mean annual increment (MAI) in trials and commercial forests of *Eucalyptus*, respectively. Similar results were found for mixture of genotypes for *Pinus* (Carter et al., 2020), where individual trees planted in mixed rows had approximately 7% greater volume compared to the ones than were planted in pure rows; and *Populus* (Foster et al., 1998), where the mixture presented 27% increase in volume over the best clone. The good performance of the mixture of genotypes can be due to a better exploitation of environmental resources for the allo-competitor genotypes. Theoretically, there is a complementarity between different genotypes when allocated in the same area as suggested by the niche partitioning hypotheses (Young, 1981). Many questions remain unexplored in this topic, such as the number of genotypes that should be planted together, i.e., the size of the CC, and the impacts of indirect genetic effects (IGE) on the performance of different CC combinations.

Linear mixed models (LMM) are routinely used to estimate breeding values or genotypic values in tree breeding (Gezan et al., 2017; Chaves et al., 2022). The LMM was extended to include IGE as a random effect for animal or plant breeding (Muir, 2005). The IGE can be described as the influence that an individual’s genes pool exerts over the phenotype of neighboring conspecifics, which can be studied by variance-component competition models (Muir, 2005). Some evidence suggests that IGE, also called associative or competition effects, are a quantitative genetic trait (Griffing, 1967), thus it can contribute to the heritable variation of a population. More information can be found in (Walsh and Lynch, 2018, Chapter 22) and (Resende et al., 2014, Chapter 11). A spatial LMM accounting for IGE, has been applied in few evaluations (Resende et al., 2005; Costa e Silva and Kerr, 2013; Herńandez et al., 2019) to forest genetics and breeding. The competition models expand the classical quantitative genetic model *P_il_*= *G_i_* + *E_il_*, in which *P_il_* is the phenotypic value of individual *i* into the environment *l*, *G_i_* is the genetic effects of individual *i*, and *E_il_* is the environmental effects individual *i* into the environment *l* by considering IGE. In this philosophy, a genotype’s total genotypic value (TGV) comprises its own genetic merit or direct genetic effects (DGE) summed to the weighted IGE that it exerts over its neighbors.

Since IGE are expected to be common in nature (Sakai, 1955), their impacts in the magnitude and direction of response to selection have been investigated by in plant and animal breeding. Muir (2005) proposed that incorporating IGE into breeding programs can increase the accuracy of selection and improve breeding outcomes. The total heritable variation that determines the potential of a population to respond to selection depends on the DGE and IGE covariance (Bijma, 2011). Costa e Silva and Kerr (2013) used simulated data to explore the impacts of different levels of genetic relatedness within the neighborhood and overall survival on the ability to estimate IGE. Costa e Silva et al. (2017) and Nunes et al. (2018) showed the implications of considering direct and indirect genetic effects on forest management and breeding programs. The complete reviews of Bijma (2011, 2014) provide more examples and details.

Based on the aforementioned, we aimed to fit a spatial competition linear mixed model for estimating genetic parameters and studying the impacts of IGE. We also propose a strategy to predict the CC with the highest potential to form highly productive commercial stands. To the best of our knowledge, no previous study has explored the prediction of CC accounting for IGE. In fact, most of them restrict inferences only to traditional linear mixed models. The main advantage of our methodology is combining genetic competition and autoregressive residual models to predict the total genotypic value (TGV) of a CC that has not yet been planted. The proposed approach was illustrated on a dataset of *Eucalyptus* breeding trials from a Brazilian wood pulp company.

## 2. Material and Methods

### 2.1. Trial description

This study was carried out in a clonally replicated trial of *Eucalyptus urophylla* × *Eucalyptus grandis* hybrids belonging to CENIBRA Celulose Nipo-Brasileira S.A. The field experiment was a randomized complete block design with 24 replications and one tree per plot, where 76 clones were planted including six commercial checks. Twenty-four copies of each clone were obtained by vegetative propagation and planted in an inter-row inter-column spacing of 2.5 x 3 m, respectively, which gives a plantation density of 1333 trees *ha^−^*^1^. The experimental area is located in the municipality of Ipaba, Minas Gerais state, Brazil, at 19° 20S, 45° 25W and 213 m of altitude. This region has two well-defined seasons: one hot and humid (October to May) and the other cold and dry (April to September) with a minimum and maximum temperature of 13.5 and 34.6 °C, respectively, and monthly rainfall ranging from 0 to 112 mm.

The mean annual increment (MAI, *m*^3^*ha^−^*^1^*year^−^*^1^) (Eq. 1) was calculated as the quotient of the volume of individual trees (*V OL_ind_*, *m*^3^) collected at 3 and 6 years after planting. This volume was extrapolated to 1 *ha* (*V OL_ha_*, *m*^3^) as shown below.

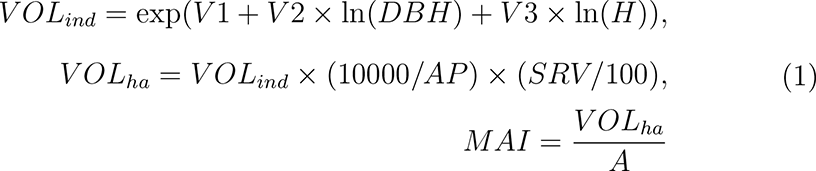

where DBH is the diameter at breast height in *cm*, H is the total height in *m*, V1 = −10.300612, V2 = 1.689752, and V3 = 1.292432. AP is the occupied area per plant, and SRV is the percentage of survival. Parameter A is the age (3 and 6) in years.

### 2.2. Statistical Analyses

In this study, we fitted three linear mixed models. The traditional linear mixed model (TM) was used in this study as a benchmark and is described below:

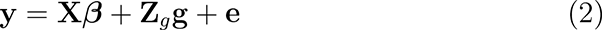

where **y** is a (*n ×* 1) vector containing the phenotypes records, which *n* the number of records; **X** is the *n × r* incidence matrix identifying which of the *r* fixed effects are associated with each observation; ***β*** is the (*r ×* 1) vector of fixed effects (intercept and 24 complete blocks); **Z***_g_* is the (*n × m*) incidence matrices relating phenotypes records to their genotypic effects contained in the (*m ×* 1) random vector **g**, with 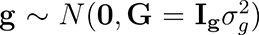 and in the (*n ×* 1) vector or random residual with 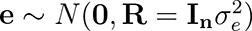. **I** is an identity matrix of proper order, 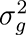 is the genotypic variance and 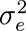 is the residual variance.

The spatial model (SM), uses a separable first-order autoregressive process in two directions (AR1 x AR1) for modeling the covariance matrix of residual effects taking an accounting of both rows and columns directions Gilmour et al. (1997):

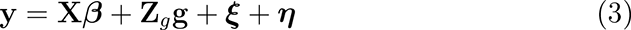

where **y** is a vector of phenotype records, taken in the field order by putting each column beneath another. The vector ***ξ*** includes random effects that represent spatial trends revealed by the correlated residual structure in which the rows and columns are autoregressive, 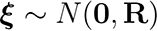, with 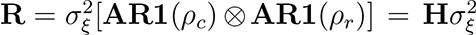. The separability property of the stochastic process ensures 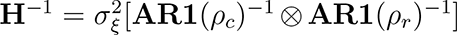 enabling the computation of **H***^−^*^1^, which is needed for estimation of variance components and for prediction of genotypic values. The quantity 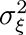 is the variance of spatially dependent residuals, **AR1**(*ρ_r_*) and **AR1**(*ρ_c_*) are the first-order autoregressive correlation matrices for 48 rows and 38 columns, respectively (Gilmour et al., 1997). The operator *⊗* represents the Kronecker product between columns and rows auto-regressive processes. The ***η*** is an (*n ×* 1) vector that encompasses independent random effects of residuals, the so-called nugget effect in the geostatistical context, representing any other influences that were not accounted in the model, where 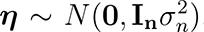, and 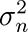 is the independent error variance. The associated mixed model equations for BLUP of genotypic values:

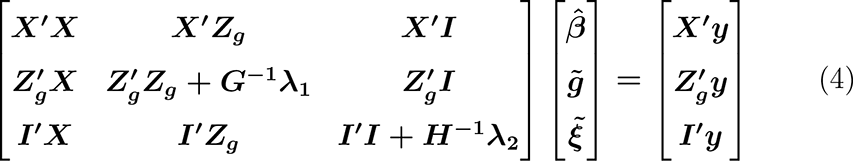

where, 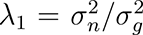, and 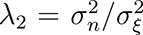. **G** and **H** are correlation matrices for genotypes (*g*) and spatial errors (*ξ*) effects, respectively.

We used Cappa and Cantet (2008) proposal, which account for the competition intensity factors (*f_ij_*) that neighbors trees *j* = *j*_1_*, j*_2_*, …j_m_* exert over a focal tree *i*, being *m* the maximum number of neighbors for a focal tree *i* (Figure 1).

**Figure 1:**
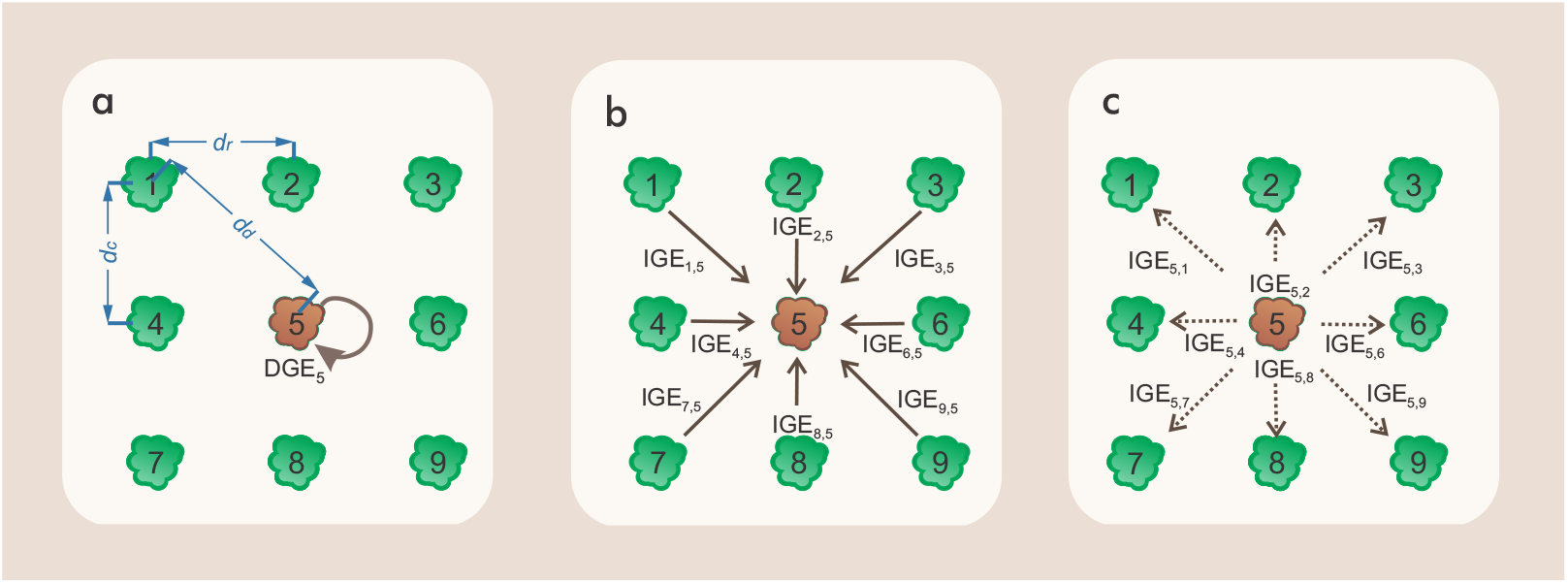
Illustration of: (a) direct genetic effect (DGE); (b) indirect genetic effects (IGE) of the *j* neighbors on the focal individual *i* ; and (c) IGE of a focal individual *i* on its *j* neighbors. The focal individual *i* stands out by a different color and is represented by the number 5; its *j* neighbors are represented by the remaining numbers (the number of neighbors can vary from 0 to *m* depending on the position of focal individual *i*); *d_r_*, *d_c_*, and *d_d_* are the distance between the focal tree and its neighbors in the row, column and diagonal, respectively.

The *f_ij_* will vary according to the distance between these individuals in the field (Figure 1a). The magnitude of *f_ij_* is related to the inverse of the distance between individual *i* and its *m* neighbors. The *f_ij_* derivations were demonstrated by Cappa and Cantet (2008) for equal inter-row and intercolumn spacing and Costa e Silva and Kerr (2013) for different inter-row and inter-column spacing. In this context, IGE can be regarded as the sum of effects that the *m* neighbors exert over the focal individual *i* 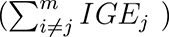 - (Figure 1b) or as the mean IGE that the focal individual *i* exert over its neighbors 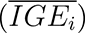 - (Figure 1c).

In matrix notation, the SCM is given by:

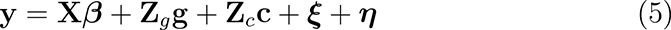

where **Z***_g_* and **Z***_c_* are (*n × m*) incidence matrices linking phenotypes records to their DGE and IGE contained in the (*m ×* 1) random vectors **g** and **c**, respectively. **Z***_g_* is a diagonal matrix with elements on the diagonal equal to 1 in the column belonging to **g***_i_* an off-diagonal equal to 0. Matrix **Z***_c_* was constructed following Costa e Silva and Kerr (2013). More details can be found in the appendix. The associated mixed model equations for DGE and IGE of clone values:

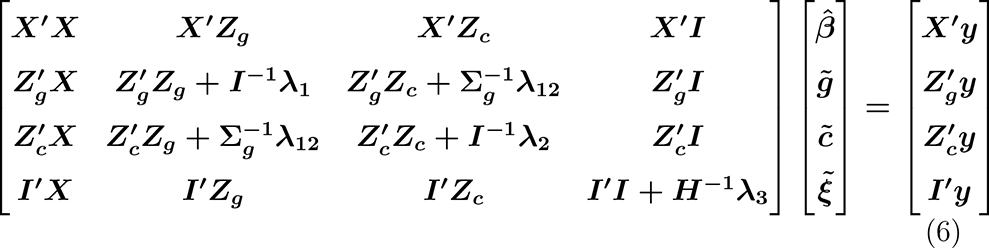

The variance-covariance matrix of the *g* and *c* (**Σ_g_**) can be given by:

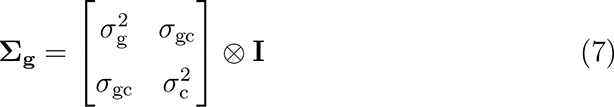

where *σ*_gc_ denote the estimates of the covariance between *g* and *c*, and **I_n_** is an identity matrix of order *n*. If *σ*_gc_ = 0, the *λ_i_* coefficients in the mixed model equations are equal to: 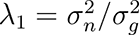, 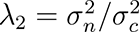, *λ*_12_ = 0 and 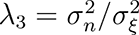 .

The total genotypic value for the focal individual *i* (*TGV_i_*) (Eq. 8) was used to rank and select individuals.

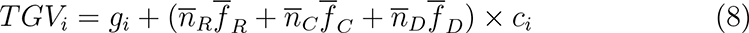

where *g_i_* and *c_i_* are the direct and indirect genetic effect, respectively; 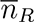, 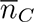, and 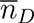 are the average number of neighbors of individual *i* in row, col, and diagonal directions, respectively; and 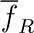, 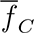, and 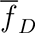 are the average *f_ij_* for row, column, and diagonal directions as proposed by (Costa e Silva and Kerr, 2013), which are represented in Eqs. 9, Eq. 10, and Eq. 11, respectively:

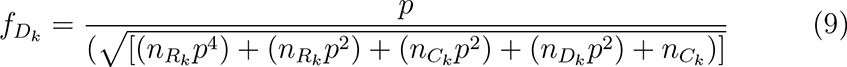

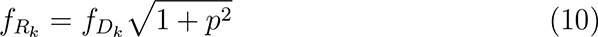

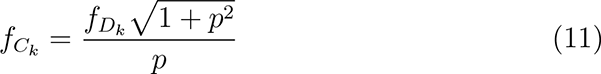

where *p* = *d_C_/d_R_*.

For clarity, the term 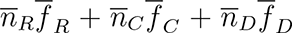 will be called competition intensity factor (CIF).

### 2.3. Model Selection and genetic parameters

The likelihood ratio test (LRT) was estimated considering an *α* = 0.005 by contrasting the likelihood of the full (*full*) and reduced model (*red*):

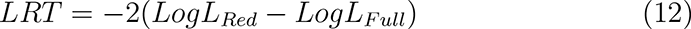

The goodness-of-fit of the evaluated models was assessed by the Akaike information criterion (AIC) (Akaike, 1974):

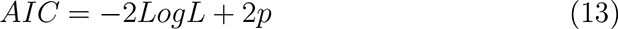

where *p* is the number of estimated parameters. The most suitable model is the one that presents the lowest values of AIC.

#### 2.3.1. Estimation of genetic and non-genetic parameters

After selecting the best fitted models among the tested ones, we estimated the broad sense heritabilities (*H*^2^) for the TM, SM, and SCM:

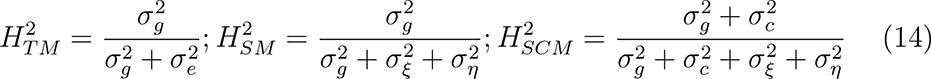

The broad sense total heritability 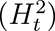 was estimated for the SCM, based on the total heritable variance 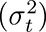as suggested by Bijma (2011) and Costa e Silva et al. (2013):

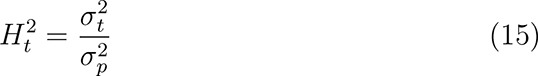

where 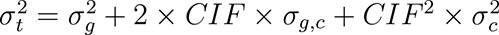

We also estimated the accuracy for DGE (*r_g_*) and the accuracy for IGE (*r_c_*) according to Mrode (2014):

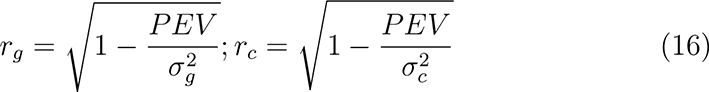

where *PEV* is the prediction error variance. The individual reliability for DGE and IGE is the square of *r_g_* and *r_c_*, respectively.

For comparison, we estimated these parameters in the traditional models and in the spatial-competition model. We compared the ranking of genotypes provided by each model using the Spearman rank correlation.

We also predicted the response to selection (*R*), in percentage, for each model, using BLUP for TM and SM and TGV for SCM, as given by:

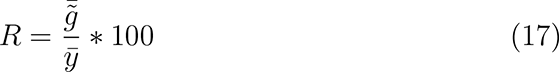

where 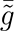 is the mean eBLUP of the selected clones (for TM and SM) or TGV (for SCM) of the selected clones using a selection intensity around 26% (20 clones out of 76), and 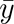 is the phenotypic mean for the evaluated trait.

All analyses were performed within the R software environment, version 4.2.1 (R Core Team, 2022). The linear mixed models were fitted using ASReml-R (version 4.1) (Butler et al., 2018). Graphs and figures were made using the ggplot2 package (Wickham, 2016).

### 2.4. Prediction of Clonal composites

Leveraging the methods mentioned above, we propose the following workflow: i) fit an individual tree competition model in real data sets; ii) estimate genetic parameters and predict the 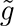, 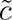, and TGV for each clone and select the best ranked ones based on their TGV; iii) investigate CCs with different composition and sizes by calculating their mean values, considering the 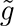 and 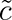 for all possible combinations of the selected genotypes for a given CC size; and iv) define a CC composition that enhances the forest productivity.

Based on the *TGV_i_* for mean annual increment (IMA), the 20 best ranked clones were selected to predict all possible combinations of clones and find the CC with the highest potential to form highly productive commercial forests. Since IGE was statistically significant, we followed our proposed workflow (Figure 2). In this study, we define the CC size as *k* = 5, since this value is used by companies in the planted forest industry. The data from the selected clones were used to create a list with all possible CC combinations. Therefore, based on the 20 selected clones, we predicted 15,504 CC combinations considering the DGE and IGE.

**Figure 2:**
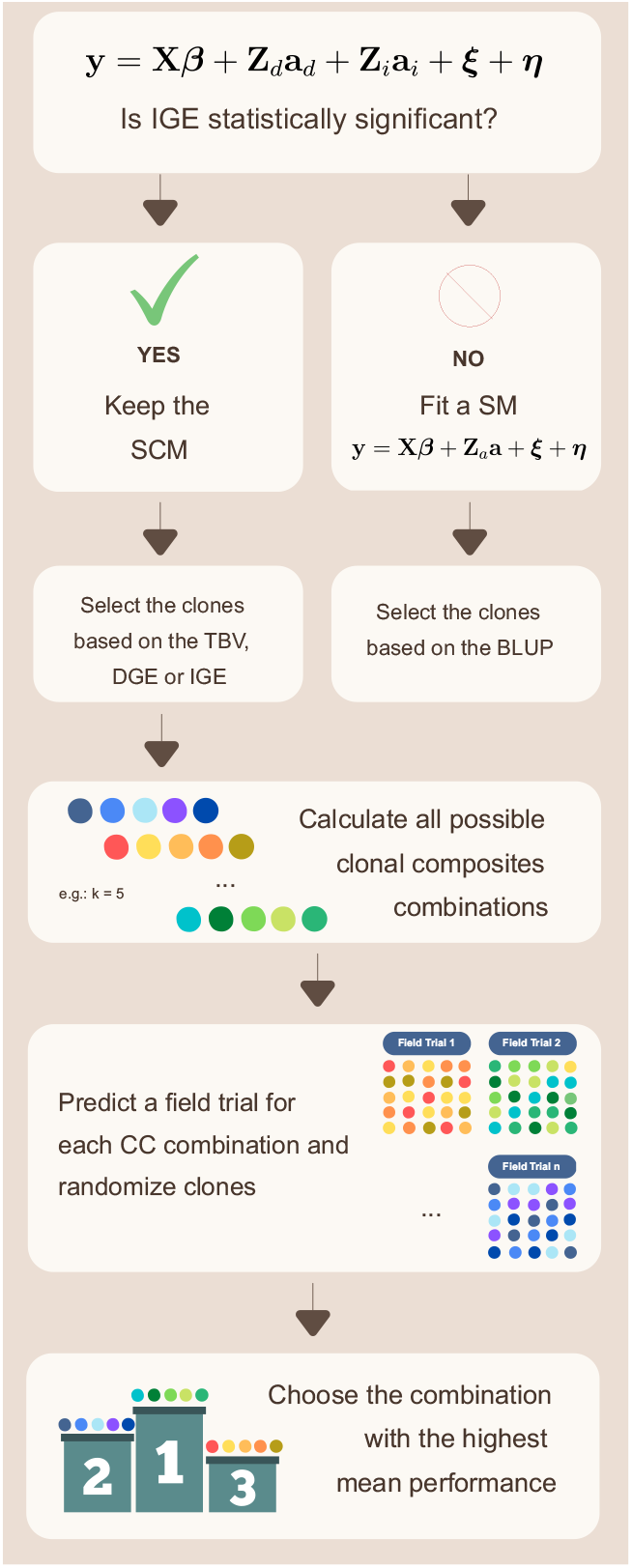
Workflow for clonal composites (CC) prediction. **y** (*n ×* 1) is a vector that contains the phenotype records, with *n* being the number of records; **X** is the *n×r* incidence matrix identifying which of the *r* fixed effects are associated with each observation; ***β*** is the *n ×* 1 vector of fixed effects, **Z***_d_* and **Z***_c_* are *n × m* incidence matrices linking phenotype records to their DGE and IGE contained in the *m ×* 1 random vectors **g** and **c**, respectively according to Silva and Kerr (2012). ***ξ*** is an effect (random) that represent spatially trends or correlated residual in the row-column structure, and ***η*** is an independent random effect of residuals, a so-called nugget effect according to Gilmour et al. (1997). TGV is the total genotypic value, DGE is the direct genetic effect, IGE is the indirect genetic effect, BLUP is the best linear unbiased prediction, SM is the spatial model, and SCM is the spatial competition model

The best CC combination was the one with the highest predicted average for our trait of interest following the (Eq. 18):

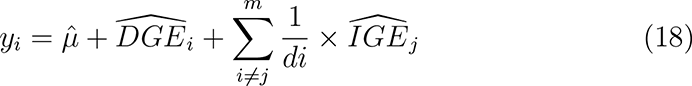

Note that the predicted mean is weighted by the inverse of the distance between rows, columns, or diagonals, which penalizes the effect of competition based on the distance between neighbor *j* and the focal individual *i* for the *m* closest neighbors. *µ* is the intercept of the model, which can be predetermined based on the mean of the field trial. The chosen CC combination is the one that presents the higher mean performance for MAI.

We propose an innovative classification for competition classes based on the magnitude of the predicted IGE and the impacts that it can have on the performance of some clones. Clones whose predicted IGE falls within one standard deviation of the mean predicted IGE were considered homeostatic, whereas clones below the threshold of one standard deviation were considered aggressive. Finally, clones above the threshold of one standard deviation were considered sensitive.

We predicted the expected mean for MAI using other strategies. In the first, we predicted a monoclonal planting by selecting the best ranked clone based on TGV. Next, we selected the five best ranked clones based on predicted DGE. Then, we selected the five most competitive clones (lowest predicted IGE). Finally, we select the three best ranked clones based on TGV and the two best ranked homeostatic clones. We also verified the effects of predicted IGE on productivity by varying the inter-row and inter-column spacing.

## 3. Results

### 3.1. Model Selection and genetic parameters

The percentages of missing observations in the trial were 13,87% and 21.93% at 3 and 6 years after planting, respectively. According to the LRT, the IGE was significant (*P <* 0.05) for both ages, suggesting that competition can arises at early ages. Thus, IGE should be accounted for when predicting the genotypic values. The correlations between DGE and IGE were negative and of high magnitude (Tables 2 and 3). For more information see Supplementary material and Figure S1. Based on the AIC, the best fitted model, for both ages, was the SCM (Table 1). This model accounts for IGE on the three directions (row, columns and diagonal).

**Table 1:** Model performance accounting for model complexity via Akaike information criterion (AIC) for a traditional model (TM), a spatial model (SM), and a spatial-competition model (SCM) accounting for the correlation between direct genetic effect (DGE) and direct genetic effect (IGE) for Eucalyptus at 3 and 6 years.

| Age |  | 3 | 6 |
| --- | --- | --- | --- |
| Model | DF | AIC | AIC |
| TM | 2 | 9503.65 | 9689.22 |
| SM | 5 | 9493.13 | 9680.43 |
| SCM | 7 | 9472.96 | 9652.06 |

DF: degree of freedom of the chi-squared distribution to perform a LRT; AIC: Akaike information criterion.

**Table 2:**
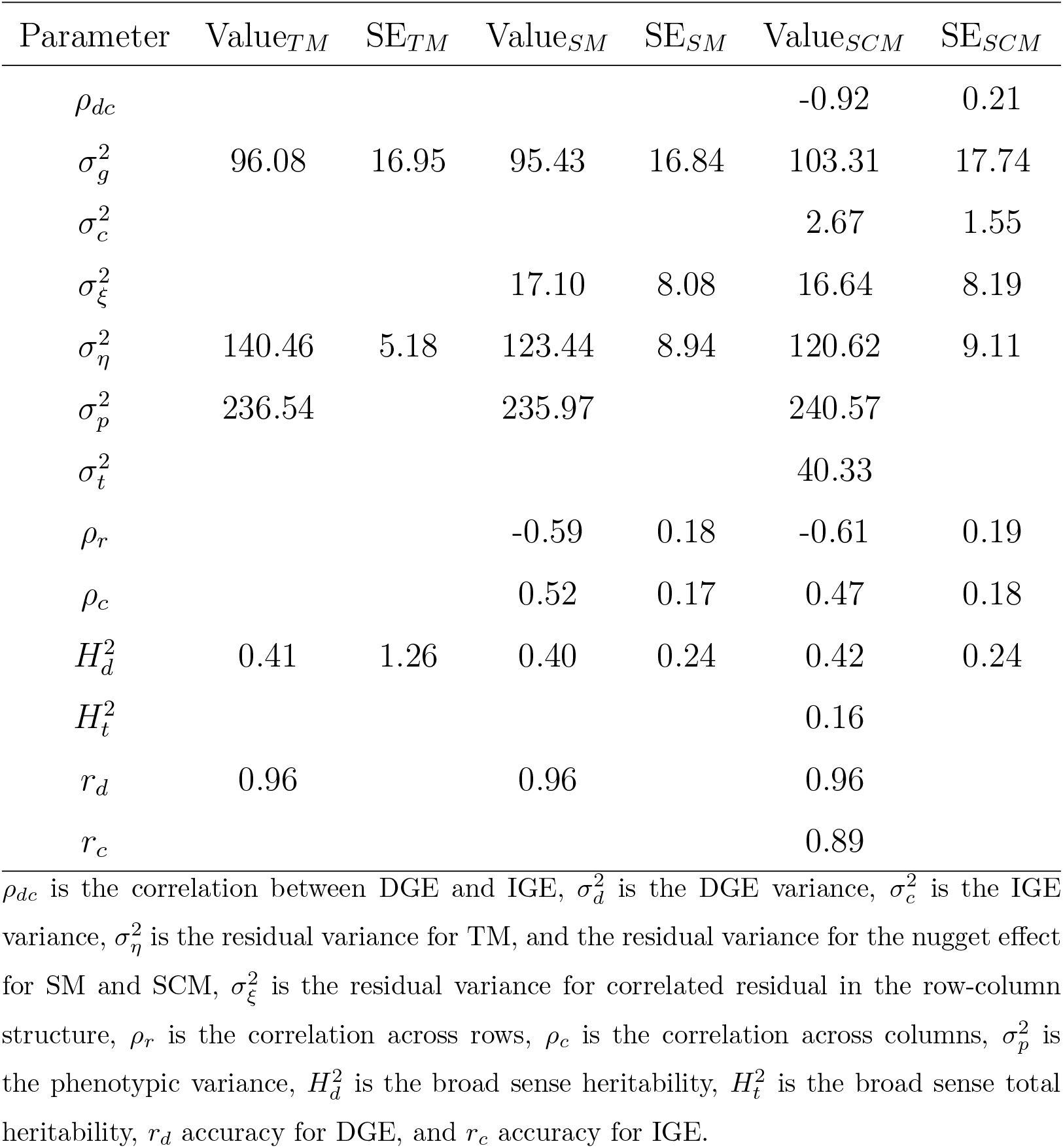
Genetic and non-genetic parameter estimates and standard errors (SE) for Eucalyptus breeding population at 3 years old by using a traditional model (TM), a spatial model (SM), and a spatial-competition model (SCM).

**Table 3:** Genetic and non-genetic parameter estimates and standard errors (SE) for *Eucalyptus* breeding population at 6 years old by a traditional model (TM), a spatial model (SM) and a spatial-competition model (SCM).

| Parameters | Value<sub>TM</sub> | SE<sub>TM</sub> | Value<sub>SM</sub> | SE<sub>SM</sub> | Value<sub>SCM</sub> | SE<sub>SCM</sub> |
| --- | --- | --- | --- | --- | --- | --- |
| $\rho_{dc}$ | | | | | -0.83 | 0.15 |
| $\sigma_g^2$ | 245.8 | 43.42 | 242.97 | 42.89 | 258.84 | 44.11 |
| $\sigma_c^2$ | | | | | 8.82 | 3.70 |
| $\sigma_\xi^2$ | | | 36.34 | 18.56 | 42.54 | 22.54 |
| $\sigma_\eta^2$ | 299.08 | 11.62 | 262.92 | 20.40 | 245.40 | 24.00 |
| $\sigma_p^2$ | 544.88 | | 542.23 | | 546.78 | |
| $\sigma_t^2$ | | | | | 111.24 | |
| $\rho_r$ | | | -0.59 | 0.19 | -0.57 | 0.20 |
| $\rho_c$ | | | 0.53 | 0.19 | 0.38 | 0.19 |
| $H_d^2$ | 0.45 | 2.89 | 0.45 | 0.56 | 0.46 | 0.80 |
| $H_t^2$ | | | | | 0.20 | |
| $r_d$ | 0.96 | | 0.96 | | 0.96 | |
| $r_c$ | | | | | 0.83 | |

$\rho_{dc}$ is the correlation between DGE and IGE, $\sigma_d^2$ is the DGE variance, $\sigma_c^2$ is the IGE variance, $\sigma_\eta^2$ is the residual variance for TM, and the residual variance for the nugget effect for SM and SCM, $\sigma_\xi^2$ is the residual variance for correlated residual in the row-column structure, $\rho_r$ is the correlation across rows, $\rho_c$ is the correlation across columns, $\sigma_p^2$ is the phenotypic variance, $H_d^2$ is the broad sense heritability, $H_t^2$ is the broad sense total heritability, $r_d$ accuracy for DGE, and $r_c$ accuracy for IGE.

The density of aggressive, homeostatic, and sensitive clones shows that most clones are within one standard deviation from the mean for both ages (Figures 3a and b). The number of different clones as neighbors that each clone had throughout the experiment (considering all replications) varied from 29 to 73, for 3 years, and from 23 to 69, for 6 years. All clones had neighbors in the 3 classes (Figure 3c and d).

**Figure 3:**
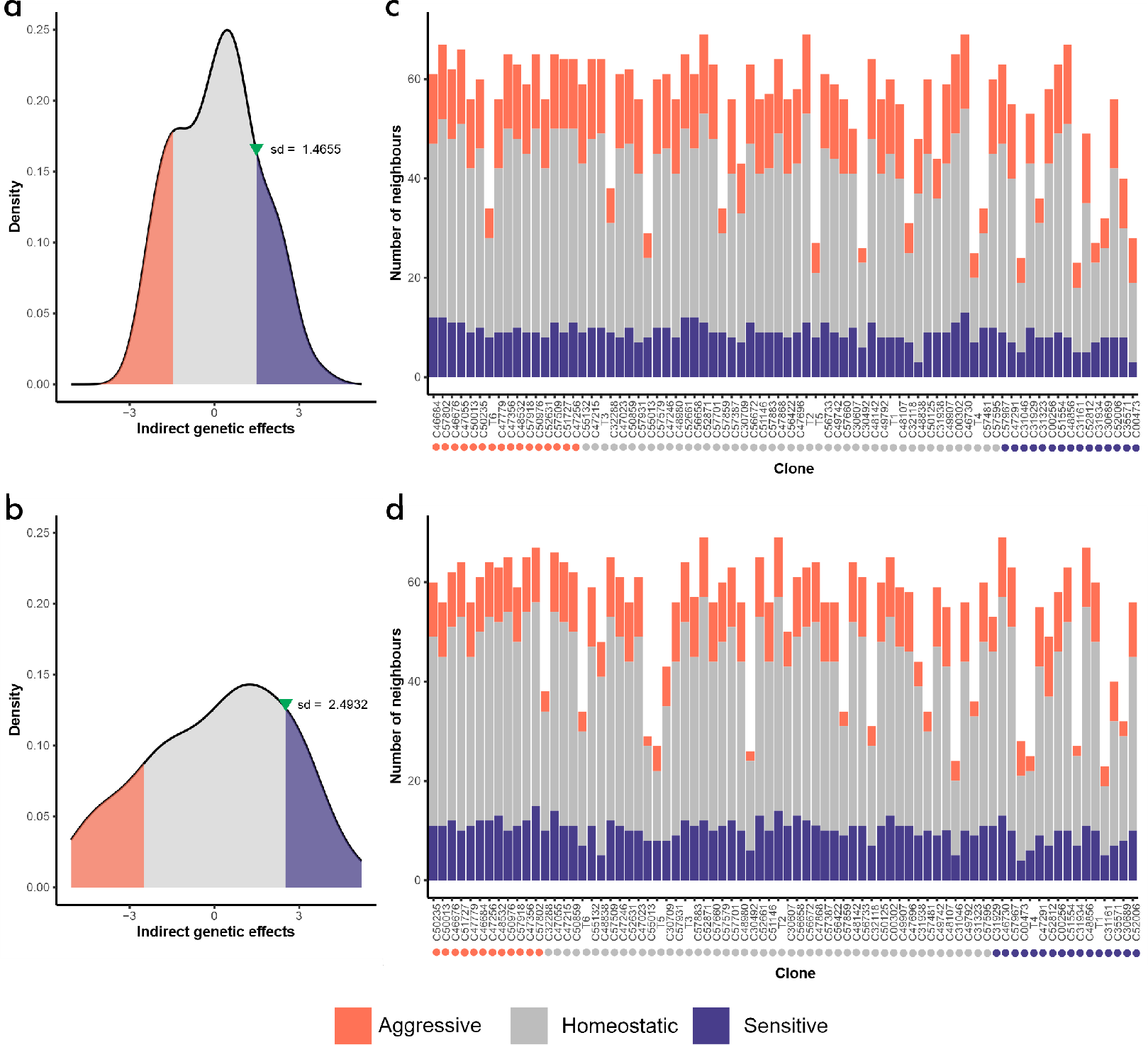
a and b) Density plots for IGE for *Eucalyptus* clones belonging to three competition classes based on the magnitude of the IGE at 3 and 6 years, respectively; Homeostatic clones are in the area between plus and minus one standard deviation from the mean IGE, aggressive clones are in the area below one standard deviation from the mean IGE (shaded orange), and sensitive clones are in the area above one standard deviation from the mean (shaded blue); The upside down green triangle indicates the point where values are higher than one standard deviation from the mean IGE; c and d) Competition class of each clone (indicated by the colored circles below the clone names) and the total number of different neighbors and their classes (indicated by the colored bars) at 3 and 6 years, respectively.

There were increments in 𝐻^2^ as the age advances (Tables 2 and 3). The heritable variation for response to selection, represented by 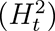 was 39% and 44% smaller using the SCM than the observed for the TM (indicated by 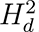), for 3 and 6 years, respectively. The proportion of the total variance explained by IGE 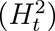 on SCM increased by 20% from ages 3 to 6 years. Resende et al. (2005) indicated that high (*>* 0.3) positive autocorrelation coefficient estimates reveal that environmental heterogeneity (spatial effects) is predominant over the competition, and negative (*< −*0.3) autocorrelation coefficient estimates indicate competition effects at the residual level probably together with environmental heterogeneity. Therefore, autocorrelation coefficients larger than 0.3 and smaller than 0.3 were used to identify dominant environmental heterogeneity and competition effects at residual level, respectively (Costa e Silva and Kerr, 2013; Cappa et al., 2015). Our results show a high level of competition across rows and high spatial environmental heterogeneity across columns (Tables 2 and 3). This probably reflects shading problems in the row direction of the field trial. When auto-correlation coefficients tend closely to 0 or 1, spatial analysis tends to give no practical improvement in the fit, despite significant changes in *LogL*. All models, for both ages, presented very high accuracy (*>* 0.95), indicating that they can be used to perform selection. The three clones that presented the lowest IGE at 3 years, meaning that they exerted the highest competition over their neighbors, were clones 16, 63, and 15. For 6 years, the lowest IGE was presented by clones 39, 37, and 15. These clones were among the ones that had the highest DGE (Supplementary material - Figure S1).

The potential of the early selection (correlation between ages 3 and 6) was assessed by the Spearman rank correlation. Lower correlations between genotypes selected at 3 and 6 years for TM and SM when compared to the SCM were found (Table 4). There were major changes in the best 20 ranked genotypes between 3 and 6 years (Supplementary material - Tables S2 and S3). The adopted selection intensity was around 26%, i.e., we selected 20 out of 76 genotypes. The proportion of the selected genotypes at 6 years that were among the previously selected genotypes at 3 years (PSG) was higher when the competition models were adopted (Table 4). For 3 years old, the response to selection by selecting the 20 best ranked genotypes were 38.29, 38.00, and 38.19% for TM, SM, and SCM, respectively. For 6 years old, the *R* by selecting the 20 best ranked genotypes were 51.16, 50.99, and 50.79% for TM, SM, and SCM, respectively. In the supplementary material, we present the DGE, IGE, their respective standard errors and reliabilities, and TGV for all evaluated clones for 3 years (Table S2) and 6 years (Table S3).

**Table 4:** Spearman rank correlation (SRC) between 3 and 6 years Eucalyptus genotypes for all evaluated genotypes (n = 76) and proportion of the selected Genotypes (PSG) at 6 years old that were among the 20 genotypes previously selected at 3 years old (n = 20).

| | TM | SM | $SCM_{TGV}$ | $SCM_{DGE}$ | $SCM_{IGE}$ |
| --- | --- | --- | --- | --- | --- |
| SRC | 0.85 | 0.80 | 0.98 | 0.99 | 0.99 |
| PSG | 0.35 | 0.35 | 0.80 | 0.90 | 0.90 |

TM: traditional linear mixed model; SM: spatial model, SCM: spatial-competition model (SCM).

### 3.2. Clonal Composite predictions

We identified aggressive, homeostatic, and sensitive clones based on the magnitude of IGE, which can help choose individuals to compose the CC. To define which ones will be the five selected clones, we use the TGV at 6 years of the 20 best ranked individuals. We predicted 15,504 CC combinations and chose the one that predicted the largest planted forest production per area accounting for the competition among clones. The chosen CC composition was the one that contains the individuals ranked in the positions 1, 2, 3, 4, and 6 for TGV; 1, 2, 4, 5, and 9 for DGE; and 2, 9, 6, 4 and 14 for IGE (sort smallest to largest). The chosen CC composition expected MAI was 58.13 *m*^3^*ha^−^*^1^*year^−^*^1^ (Table 5). Despite C47055 not being ranked among the top five clones for TGV and DGE (Table 5), it was selected based on the adopted strategy. The IGE that C47055 exerts over its neighbors is less than half the IGE that the best ranked clone (C50013) exerts over its neighbors (Table 5). Based on these results, we can infer that at least 1 out of 5 clones must be selected prioritizing the trade-off between DGE and IGE.

**Table 5:** Expected mean for mean annual increment at age 6 years for the best clonal composite composition for *Eucalyptus grandis* x *Eucalyptus urophylla* hybrids.

| Clone | DGE | IGE | MAI | $Rank_{TGV}$ | $Rank_{DGE}$ | $Rank_{IGE}$ |
| --- | --- | --- | --- | --- | --- | --- |
| C50013 | 34.80 | -4.56 | 65.08 | 1 | 1 | 2 |
| C50976 | 28.63 | -3.49 | 58.93 | 2 | 2 | 9 |
| C46684 | 27.77 | -4.22 | 58.06 | 3 | 4 | 6 |
| C51727 | 26.85 | -4.25 | 57.15 | 4 | 5 | 4 |
| C47055 | 21.17 | -2.00 | 51.43 | 6 | 9 | 14 |
| Mean | 27.84 | -3.70 | 58.13 |  |  |  |

DGE is the direct genetic effect, IGE is the indirect genetic effect, and MAI is the mean annual increment. $Rank_{DGE}$ , $Rank_{IGE}$ , and $Rank_{TGV}$ is ranking for each of the select genotypes based on the TGV, DGE, and IGE (sort smallest to largest).

Using the five most positive DGE and the five most aggressive clones, the predicted MAI were 56.35 and 55.11 *m*^3^*ha^−^*^1^*year^−^*^1^. The use of the three best ranked clones for DGE, and the two best ranked homeostatic clones provided 52.25 *m*^3^*ha^−^*^1^*year^−^*^1^. Predictions for a monoculture with the best ranked clone based on TGV provided an expected MAI of 61.29 *m*^3^*ha^−^*^1^*year^−^*^1^. Therefore, for composing a clonal composite of size five, our strategy allowed a higher mean for MAI (58.13 *m*^3^*ha^−^*^1^*year^−^*^1^) than all the other tested strategies and a slightly inferior mean compared to the monoclonal planting. The mean for all 15,504 CC combinations of size five is displayed in the supplementary Material - Figure S2.

## 4. Discussion

Our results shed new light on the potential of spatial-competition models to predict the performance of a mixture of genotypes, including CC, as a combination of direct and competition effects. This section summarises the findings and contributions made by the present study focusing on our two main goals: fit a spatial-competition model to estimate genetic parameters and investigate the effects of IGE (Section 4.1), and propose a new approach to predict CC forest production (Section 4.2).

### 4.1. Model Selection and genetic parameters estimates

In this study, we choose to work with MAI (*m*^3^*ha^−^*^1^*year^−^*^1^) because it depends on diameter at breast height and plant height. Additionally, MAI is a function of the planted area and the age of the forest. The MAI increases with the age of the stand until competition and physiological maturity slow the growth rate of the cohabitating trees. The chosen trial, established with single-tree plots, was efficient for estimating the IGE since the focal clone is often surrounded by genetically distinct others, favoring the inter-clonal competition experienced in CC plantations.

The plastic response of a given clone under competition is closely related to its ability to capture limited resources and use them efficiently in this shared environment. In the early stages, trees mainly compete for underground resources such as water and nutrients. After canopy closure, the competition for light intensifies. Therefore, competition can be considered a force that disproportionately affects cohabiting individuals’ growth. Clones present different magnitudes of IGE as age advances, indicating that a differential gene expression and, consequently, a differential gene interaction between the focal tree and its neighbors can occur. Therefore, the 8.06% increase in mortality observed between 3 and 6 years might be due to eco-physiological effects not accounted for by our model.

There are multiple ways to construct the **Z_c_** matrix and calculate the CIF. However, the results must make biological sense. Our **Z_c_**matrix was constructed based on the CIF proposed by Costa e Silva and Kerr (2013), an extension of Cappa and Cantet (2008). It provided IGE values much lower than the DGE, which is expected since the clone’s performance is mainly affected by the pure expression of their genes due to DGE. Also, we worked with many interacting individuals, which can reduce the model’s ability to capture IGE (Bijma, 2011). The adopted modeling approach was efficient in capturing the IGE, allowing the estimation of variance components and predicting TGV on the individual level. Also, the classification of clones in competition class was a refined way of defining classes based on a common statistical concept. Most clones were homeostatic, which may lead to more phenotypic stability when planted in marginal areas.

The notion of genetic architecture and the response to selection can be altered by accounting for IGE. IGE can amplify or diminish the response to artificial selection compared to what is estimated via BLUP based heritability (Costa e Silva et al., 2013). The range of heritability values depends on the covariance between DGE and IGE, which is directly affected by the genetic composition, presence of inter-genotype competition, density, and spacing. We observed a strong and negative covariance between DGE and IGE, which impact the heritable variation, decreasing the potential response to selection for MAI based on individual performance. Bijma (2011) speculates that this might be due to evolutionary mechanisms that selected related genotypes in natural populations with good growth capacity. As a result, there is a less heritable variation to be explored by accounting IGE. Thus, the predictions in response to selection may be biased due to ignoring the IGE when it is statistically significant. The opposite can occur when the covariance between the direct and indirect effect is positive, as evidenced by Costa e Silva et al. (2013) for resistance to *Mycosphaerella* leaf disease. Future studies can help to elucidate the physiological aspects of competition, including how stress signals are perceived and how they trigger genetic and metabolic responses.

### 4.2. Clonal Composite predictions

Whether or not IGE is considered can significantly alter the ranking based on TGV. The predictions for monoclonal fields using the best ranked clone remains the most productive. This result corroborates with the response to the selection philosophy that expects the highest genetic gain when selecting the highest performance genotype. Therefore, we believe that CC should be planted when the goal is reducing the risks associated with growing pure stands of a single clone.

Competition models can provide valuable information to predict CC composition, since changes in response to selection and ranking may occur by accounting for the IGE. Methods that use a spatial-competition model for predicting CC performance were yet to be implemented. The magnitude and direction of IGEs may vary depending on the size and composition of the CC, the availability of resources, and the number of neighbors. At this stage, our discussion was guided by the following questions: i) what is the ideal size of the CC? ii) should we mix competitive and non-competitive clones into a CC? and iii) are the best clones for DGE the best for composing the CC?

Clones that are good competitors perform better due to the IGE that they exert over their neighbors, which can increase their 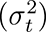. Therefore, the ideal size of the CC will be based on the researcher’s previous experiences, which must consider: the maintenance of a reasonable level of diversity, the availability of seedlings or seeds, and the trade-off between DGE and IGE. The use of homeostatic clones associated with the best ranked TGV provided a reasonable performance compared to the best CC, suggesting that it can be explored in situations of denser spacing or longer cutting cycles (where a greater magnitude of IGE is expected) without sudden decreases in productivity. The best ranked genotypes will not always be the most appropriate to compose a CC due to the trade-off between DGE and IGE. However, the proportion of each clone into the CC is not yet optimized as it considers the same contribution proportion (20%) for each of the five clones. More elaborated solutions can be explored by optimization CC via linear programming techniques.

Our strategy should be explored to predict the performance of clonal composites that have been untested in the field. Testing all possible combinations of clones to define a compound is costly, time-consuming, and, in some situations, impractical. We successfully fit parsimonious spatial-competition models, extracting useful information to predict many CC combinations and their expected average performance. We identified genotypes with distinct competition response patterns using the IGE magnitude, which can be a piece of valuable information for assisting with strategic decision-making within tree breeding programs or commercial operations. For example, we can predict the expected changes in a trait performance due to modifications in the number of clones composing the CC or their phenotypic plasticity. Clearly, the SCM model provided more information to assist the selection process than the TM and SM models. Thus, the SCM model can be expanded to different situations of agriculture and plant breeding, helping to answer important and under explored questions regarding management and mixture of genotypes.

## 5. Conclusion

We showed based on a spatial-competition model that overestimated heritability is obtained when indirect genetic effects are not considered for mean annual increment in tropical eucalyptus. Also, a methodology for predicting high-performance clonal composites based on the trade-off between direct and indirect genetic effects was proposed. Therefore, predicting clonal composites by capitalizing on the indirect genetic effects can generate a strategic advantage in determining the best clonal composite combinations to be planted.

## 6. Author contribution statement

Ferreira, F.M.; and Dias, K.O.G., designed the research. Ferreira, F.M.; Chaves, S.F.S.; and Dias, K.O.G., performed the statistical analyses and wrote the first draft. Bhering, L.L; Alves, R.S; Resende, M.D.V; Gezan, S.A; Viana, M.S; and Fernandes S.B. revised drafts of the paper. Alves, R.S; Takahashi, E.K; and Souza J.E. provided the *Eucalyptus* dataset. All of the authors read and approved the final manuscript.

## 7. Supplementary material

The Supplementary Material is available at: https://github.com/filipe-manoel/ me/blob/master/Supplementary_Mat_IGE_CC.pdf.

## Acknowledgements

This work was financially supported by the Conselho Nacional de Desenvolvimento Científico e Tecnológico (CNPq) and Coordenação de Aperfeiçoamento de Pessoal de Ńıvel Superior (CAPES) – Finance Code 001. The authors also acknowledge the company CENIBRA Celulose Nipo-Brasileira S.A for the *Eucalyptus* data and all the support.

## Appendix

Bellow we have a small example of how to construct the incidence matrices **X**, **Z_g_**, and **Z_c_** for a complete randomized block design with 6 treatments and two blocks and one plant per plot (Figure 4). The first block is represented in the left and the second block in the right. The plots from 1 to 6 are from block 1 and the plots from 7 to 12 are from block 2. The hypothetical distance between the focal tree and its neighbors in the row, column and diagonal are 2.5, 3, and 3.90 m, respectively.

**Figure 4:**
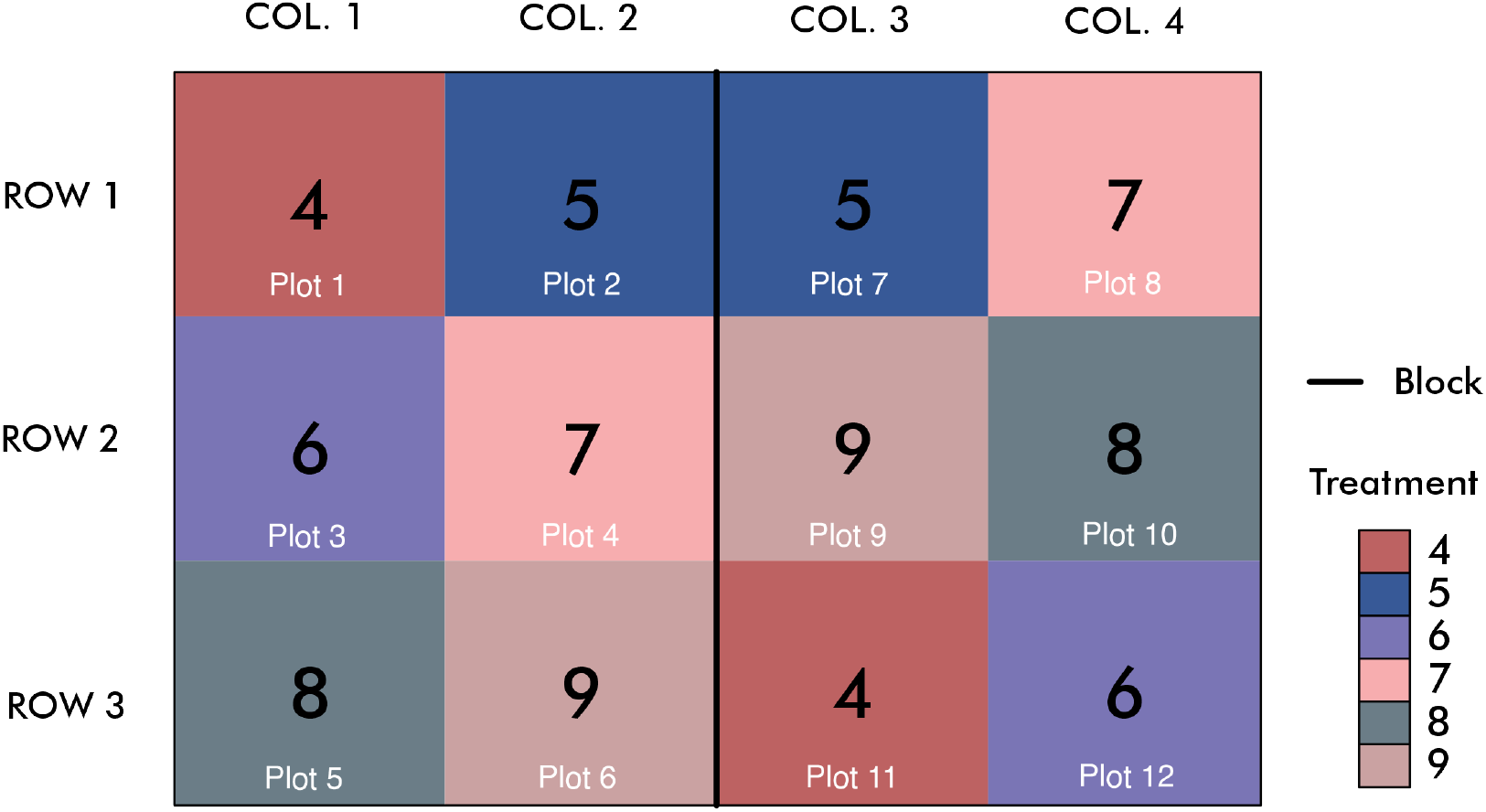
Complete randomized block design for a breeding trial with 6 treatments and two blocks and one plant per plot. The colors represent a production for an a hypothetical trait.

We wish to fit the SCM (Equation 5) to this data. Therefore, our incidence matrix **X** can be written as:

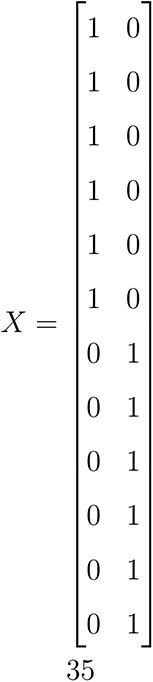

The **Z_g_** can be written as:

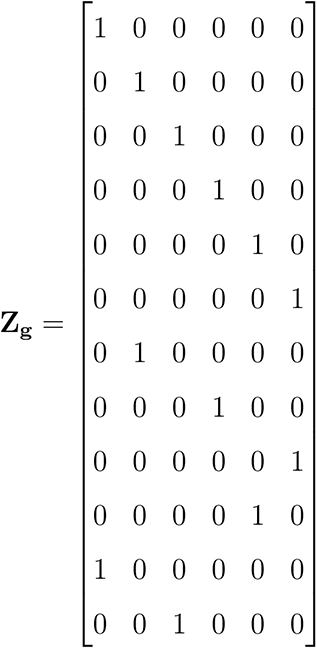

Finally, the **Z_c_** can be written using the competition intensity factors (CIF) by applying the equations 9, 10, and 11.

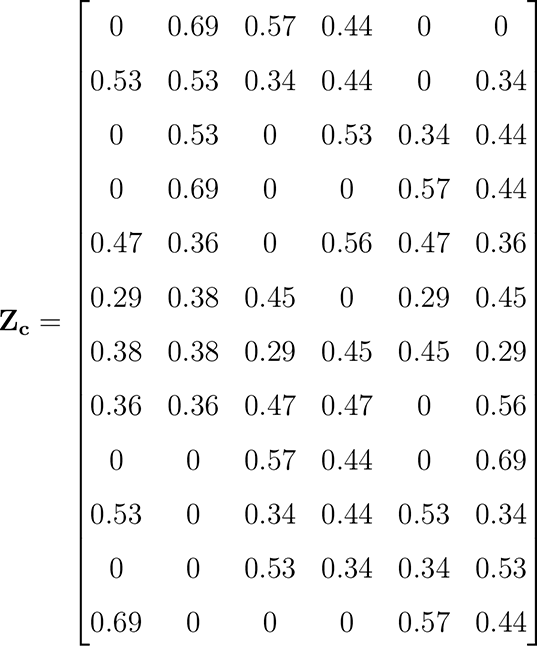

